# Density-Dependent Color Scanning Electron Microscopy (DDC-SEM) for calcified tissue and pathological calcification

**DOI:** 10.1101/2022.05.06.490880

**Authors:** Elena Tsolaki, Adrian H Chester, Sergio Bertazzo

## Abstract

Scanning electron microscopy (SEM) is widely used for materials characterization. It has also been successfully applied to the imaging of biological samples, providing invaluable insights into the topography, morphology and composition of biological structures, including pathological minerals, in diseases affecting cardiovascular, kidney and ocular tissues. Here we provide a comprehensive and detailed guide on how to use colored SEM to aid the visualization and characterization of pathological calcification, and identify the effects of different sample preparation protocols for the visualisation of these minerals.

---

Pathological calcification, or the formation of minerals in different tissues, is a process well-known as it has been observed in a number of diseases, including several cancer types; including breast, ovary and prostate cancer [1-3], tuberculosis [4], chronic kidney disease [5], aortic valve stenosis[6] and rheumatic heart disease [7]. Among the major challenges in the study of pathologic calcification are difficulties in the identification and visualization of minerals [8-12]. This is due to their nanometric scale, and the fact that they are not easily distinguished within the organic matrix [8-12].

Scanning electron microscopy (SEM) has been commonly employed most widely to the study of calcium phosphate minerals [8-15] and a method using a combination of different SEM detectors has proven to be particularly useful for the identification and visualization of the minerals present in cardiovascular, kidney and ocular diseases [8-14]. The method (named density-dependent colored SEM (DDC-SEM)), consists of manipulation of the original SEM images to produce colored images that provide both topographic and density information (associated to the atomic number of the elements present in the sample (Fig.1)) without any staining required [10,16,17].

**Figure 1.**
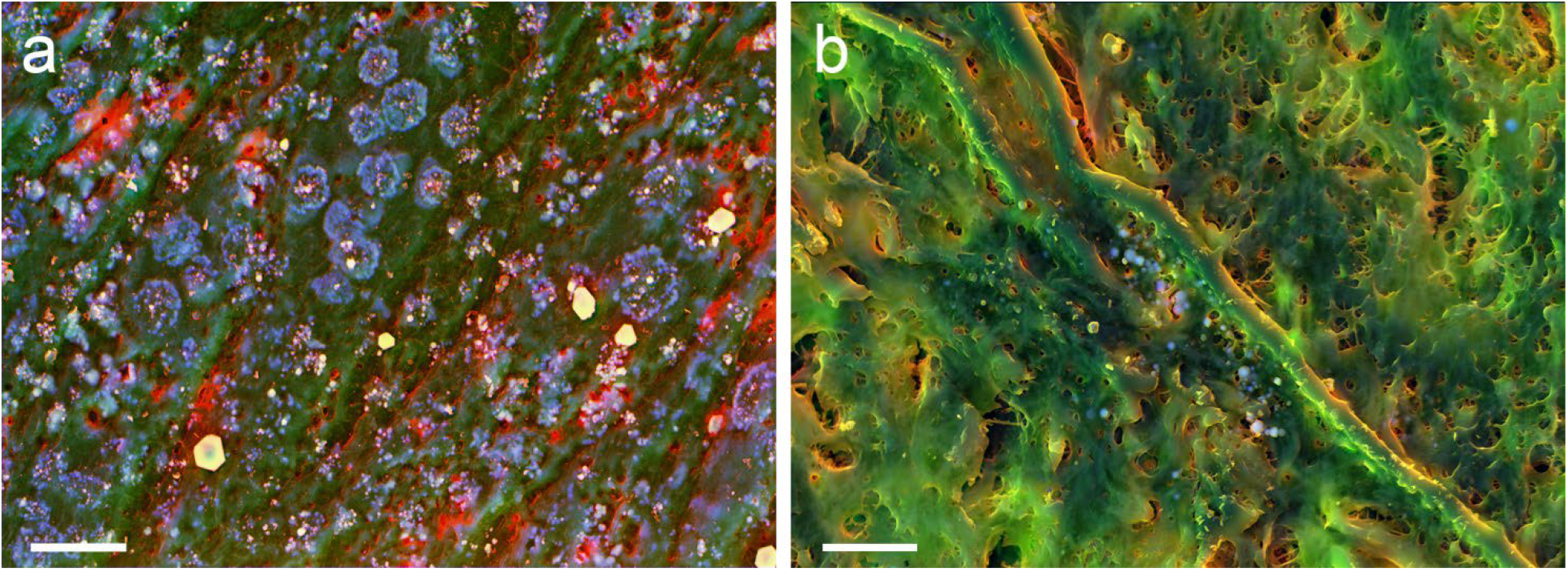
DDC-SEM micrographs of different sample types, where organic and inorganic material can be distinguished based on their colour. (**a**) DDC-SEM of vascular smooth muscle cells, with blue and yellow being the inorganic, and green and red the organic material. Scale bar = 10 µm. (**b**) DDC-SEM of a histological slides of cardiovascular tissue, with green and red being the inorganic material and blue the organic material. Scale bar = 5 µm.

The DDC-SEM approach has been first suggested over a decade ago [16-18], and was initially used to add color to the otherwise black and white SEM images, for aesthetic purposes. Recently, its potential for the study of pathological calcification has been rediscovered [8-12]. The DDC-SEM method allows for a clear visualization of the organic and inorganic material present in all types of soft tissues, owing to the contrast generated by the substantial differences in atomic number and density between the soft tissue and the mineral present in it.

Here we describe in detail sample preparation and computational practices methods to obtain high quality DDC-SEM images from pathological calcification samples. The suggested protocols allows for clear visualization of the organic and inorganic material present in all types of soft tissues, owing to the contrast generated by the substantial differences in atomic number and density between the soft tissue and the mineral present in it. These methods can be applied to all types of pathological calcification observed in any soft tissue and also to mineralizing in vitro models.

Human vascular smooth muscle cells (HVSMC) were acquired from ATCC®. The cells used were kept at -80°C and were used at passage four to eight. Cells were cultivated in vascular cell basal medium (ATCC®) complemented with vascular smooth muscle growth kit (ATCC®).

For the DDC-SEM methods to be applied to cell mineralizing in vitro models, the following protocol was found to give the best results:

1. Cells should be fixed in their wells in a 4% (mg/ml) formaldehyde in phosphate buffered saline solution of a composition of 0.137M sodium chloride, 0.0027M potassium chloride, 0.01M sodium phosphate dibasic and 0.0018M monopotassium phosphate (pH 7.4) at 4°C, for at least 24 hours.
2. Samples must be washed in water three times for five minutes.
3. Samples must then be dehydrated through a series of graded ethanol solutions; 20%, 30%, 40%, 50%, 60%, 70%, 80%, and 90% for a 10-minute interval each, followed by three changes of pure ethanol for a 10-minute interval each.
4. The samples should then be air dried.
5. If cultured in well plates, the bottom of the wells should be carefully punched out using a steel hole punch tool of appropriate size.
6. Each of the samples must then be secured onto an aluminium stub using conductive carbon adhesive tape, carbon coated with a 10nm layer, and silver painted thoroughly, from the edges of the sample to the aluminium stub.

Tissue samples were obtained by the Old Brompton Hospital-London and by the Oxford Heart Valve Bank at John Radcliffe Hospital – Oxford, the ethical approval for which was obtained for the original research paper [10].

Biopsies of a few centimetres were randomly obtained from tissues, which can be sectioned into small samples of two to five millimetres. With the samples sectioned, the following protocol should be applied:

Samples must be fixed in a 4% (mg/ml) formaldehyde in phosphate buffered saline solution of a composition of 0.137M sodium chloride, 0.0027M potassium chloride, 0.01M sodium phosphate dibasic and 0.0018M monopotassium phosphate (pH 7.4) at 4°C, for at least 24 hours.

1. Samples must then be dehydrated through a series of graded ethanol solutions; 20%, 30%, 40%, 50%, 60%, 70%, 80%, and 90% for a 10-minute interval each, followed by three changes of pure ethanol for 10-minute each.
2. The samples should then be left to air dry.
3. Each of the samples must then be secured onto an aluminium stub using conductive carbon adhesive tape, carbon coated with a 10nm layer, and silver painted thoroughly from the edges of the sample to the aluminium stub.

Paraffin embedded histological slides of four to ten micrometres can be used alternatively. The paraffin wax must be removed by immersing the slide in pure xylene twice, for 10 minutes each time, before the protocol described above is applied.

It is crucial that samples should be silver painted thoroughly from the area surrounding the tissue to the silver stub. This step it is likely the factor determining the quality of the images to be obtained in the SEM.

For DDC-SEM, samples should be imaged in a SEM equipped with at least secondary (SE), and backscattered electron (BSE) detectors. The latest SEM microscopes may also present, for instance, an immersion lens detector, which is merely a variation on the SE and BSE detectors. Other detectors may be used to obtain different colors in the DDC-SEM images.

A voltage commonly used for imaging is 10 kV or higher. Scan speed decisions, however, require trading-off speed of image acquisition against quality of the image.

Images should be acquired as routinely for the equipment in use. The images captured should be superimposed using Image J or Adobe Photoshop CC 2019.

One micrograph can be acquired using the SE detector (or any variation of this, such as an in-lens detector, Fig. 2(a)) and another using a BSE detector (Fig. 2(b). These will then be changed into two color images, as required for the DDC-SEM:

**Figure 2.**
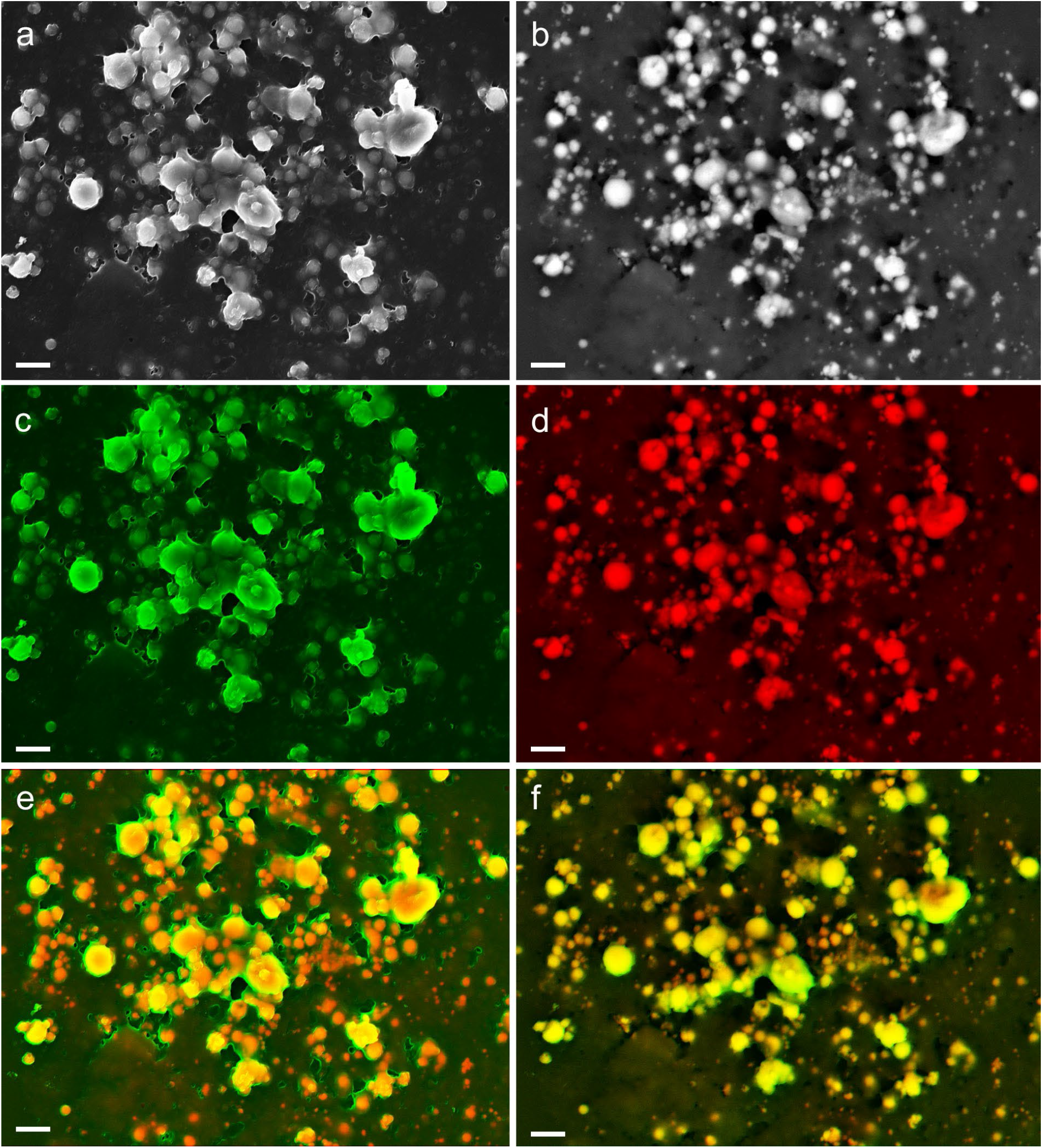
Density-dependent colored electron micrographs of calcification in a mineralizing vascular smooth muscle cell culture model. (**a**) Original in-lens detector micrograph, providing topographic information. (**b**) BSE detector micrograph, showing dense material (minerals) in white, and organic material in black. (**c**) In-lens detector micrograph assigned the green channel. (**d**) BSE detector micrograph assigned the red channel. (**e**) Superimposed DDC-SEM image, with red and orange colors indicating the calcification and green indicating the organic matrix, whilst also providing topographic information. (**f**) Similar superimposed density-dependent colored image, using the secondary detector image as the green channel and the backscattered image as the red channel. Scale bars = 2 µm.

1. A color channel should be assigned to the SE micrograph (in the example shown, the color green is assigned to the in-lens image, Fig. 2(c)) and a different color channel to the BSE micrograph (in this same case red was assigned, Fig.2(d)).
2. The two images then must be superimposed.

The resulting superimposed image will display the inorganic material in one color (red/orange) in clear contrast with the organic material, which will appear in a different color (green, Fig. 2(e)). A similar result can also be obtained using other detectors (Fig. 2(f)).

Incidentally, one of the best results we have obtained was achieved by the combination in-lens/BSE (Fig. 2f), where the topographic information on the material surrounding the mineral present in the sample (in this case, on collapsed cells and cell well) is clearer. This occurs due to the fact that the in-lens detector acts as a filter for specific electron energies, providing, in this way, the information of a smaller sample area in comparison to the traditional SE detector. Additionally, given the similarities between the contrast levels of the BSE and SE images, there is no strong color variation between the structures observed in the individual original images.

On the other hand, it is possible to take advantage of having several different detectors in the SEM. With more than two detectors, one may obtain multi colored DDC-SEM micrographs. For instance, the green, red and blue channels may each be assigned to a different detector (in-lens, Fig. 3a, SE, Fig. 3b and backscattered, Fig. 3c). The resulting image in our example (Fig. 3d) presents the inorganic material as pink and purple, as a result of the superimposed red and blue from the original BSE and SE images, whereas the organic material is seen as dark blue and green, as a result of the combination of the three original images (Fig. 3d). It is important to note that in all of these methods, the minerals and the organic material always present a consistent colour throughout the sample (this could be affected by issues in the process of imaging).

**Figure 3.**
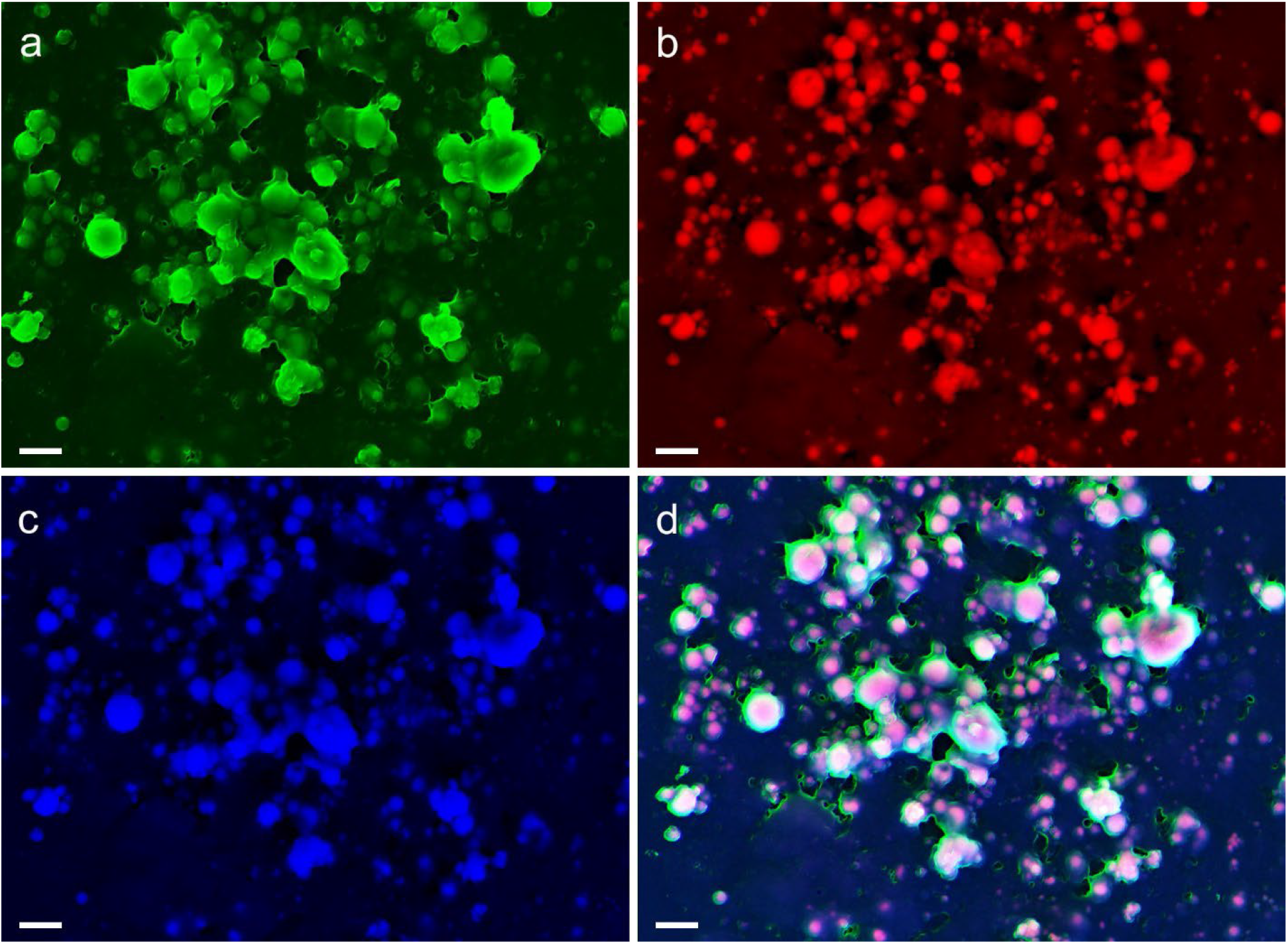
Density-dependent colored electron micrographs of calcification in a mineralizing vascular smooth muscle cell culture model. (**a**) In-lens detector micrograph assigned the green channel. (**b**) BSE detector micrograph assigned the red channel. (**c**) SE detector micrograph assigned the blue channel. (**d**) Resulting superimposed DDC-SEM image, where pink and purple colors indicate the calcification, while blue and green colors indicate the organic matrix. Scale bars = 2 µm.

In order to obtain the high quality DDC-SEM images, it is fundamental that the samples do not present any amount of build-up electrons on the surface (Fig. 4a). These could distort the image, through the formation of horizontal lines and very bright areas (Fig. 4a), which could also interfere in the color assignment to the right location in the image (Fig. 4b).

**Figure 4.**
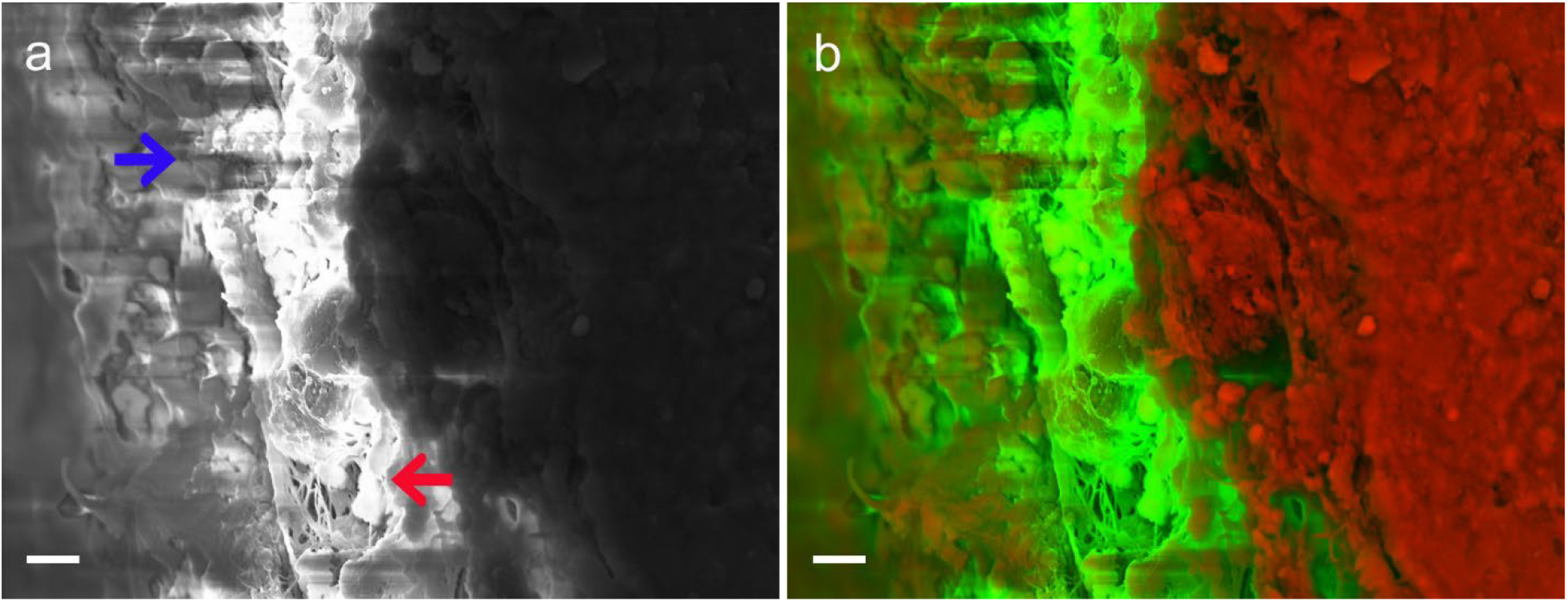
Images of inappropriately prepared aortic valve tissue. (**a**) In-lens detector micrograph of a sample where no silver paint was applied, showing the ‘charging’ effects on the sample; horizontal lines (blue arrow) and bright regions (red arrow). (**b**) Poor quality DDC-SEM micrograph of the same sample, where charging is clearly observed. Scale bars = 4 µm. Human aortic valve tissue was collected under the guidelines of ethical approval for the original study [10]. Biopsies of a few centimeters were obtained through the tissues, the samples were then cut into small pieces of two-five millimeters prior to fixation, for better fixative infiltration.

To avoid the charging effects, it is essential that the sample to be imaged is as small as can be, and, perhaps most importantly, that the sample is well secured on a metallic stub, with abundant silver or carbon paste. The conductive paste will guarantee that all electrons have a clear path to be discharged from the sample. This step is even more fundamental than the coating of the sample with a conductive layer, for which it is best to use double coating, i.e. with carbon and then with a layer of metal as thin as possible (to avoid having the metal add to the density of the whole sample). Even if still essential, the coating layer alone does not have as clear an effect on sample charging as the abundant use of a conductive paste.

Another cause of errors is the choice of inappropriate dehydration and drying methods. The most common approaches for dehydration are critical point drying [28,29] followed by the hexamethyldisilazane (HMDS) dehydration [28,30]. These methods have been developed to maintain, as much as possible, the original structures from biological samples as they would be found in the hydrated state. Air drying is, nonetheless, more suitable for the DDC-SEM to allow the imaging of minerals surrounded by organic material.

A comparison of air-dried samples and hexamethyldisilazane (HMDS) dried samples shows quite clearly that HMDS allows for greater detail of the microstructure in both the in-lens (Fig. 5(a)) and the BSE (Fig. 5(c)) images. On the other hand, it is hard to distinguish whether the structures observed in these images are organic or inorganic. Additionally, only minerals on the surface of the sample can be identified. For the air-dried sample, the organic matrix collapses (Fig. 5b), allowing for the minerals to be easily visualized, even by the in-lens detector (Fig. 5b), and minerals under a thin layer of organic material can also be observed (Fig. 5d). A comparison between the DDC-SEM micrographs produced for the HMDS prepared samples (Fig. 5e) and the air-dried samples (Fig. 5f) clearly shows that, due to poor contrast differences, it is harder to distinguish between minerals and organic material on the HMDS samples. In this case, the DDC-SEM might be misinterpreted, since some organic material falsely appears in the same color as the mineral (Fig. 5e).

**Figure 5.**
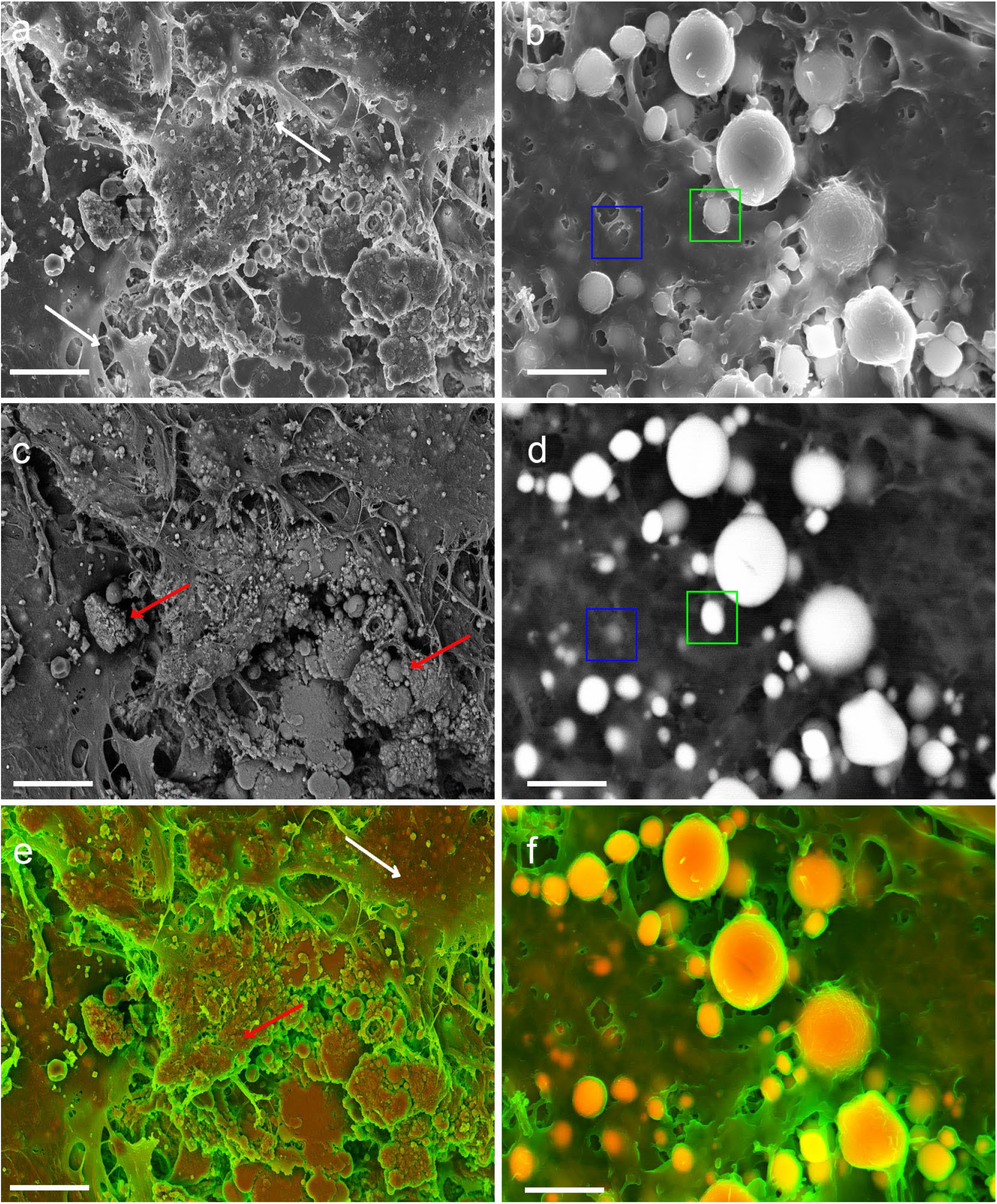
Electron micrographs of human aortic valve tissue. (**a**) In-lens detector micrograph of a HMDS dried sample, where the organic microstructure can be visualised (white arrows). (**b**) In-lens detector micrograph of air-dried sample, where the organic material has collapsed, allowing for calcification to be more easily visualised. Scale bar = 2 µm. (**c**) BSE micrograph of the area shown in a, where some calcification can be observed on the surface of the sample (red arrows). Scale bar = 4 µm. (**d**) BSE micrograph of the area shown in b, where calcification can be easily visualised with marked contrast differences between the organic and inorganic material. Calcifications that were either closer to the surface (green square) or deeper in the tissue can be equally well identified (blue square). Scale bar= 2 µm. (**e**) DDC-SEM micrograph produced by assigning the green channel to the image in a and the red channel to the image in c, with the poor contrast of the BSE micrograph resulting in an image where it is hard to distinguish between the organic (white arrow) and inorganic (red arrow) material based on color. Scale bar = 4 µm. (**f**) DDC-SEM micrograph produced by assigning the green channel to the image in b and the red channel to the image in d, where the organic and inorganic materials can be easily distinguished based on color. Scale bar = 2 µm.

These comparisons reveal the importance of air-drying for imaging of pathological calcification. Unlike other SEM biological applications, where the microstructure of the organic matrix is of interest, the collapsing of the surrounding matrix is essential for minerals to be clearly observed in soft tissues. Therefore, methods such as HMDS and critical point drying are best avoided, as they can result in misinterpretation of inorganic as organic material in backscattered images.

The use of DDC-SEM can contribute significant advances in the research field of pathological calcification, as it allows for a clear visualization of the minerals present in the samples. As regards the protocol described here, we would like to stress that inadequate sample preparation can lead to misinterpretation of minerals for organic material, and reiterate that air drying has been identified as the best sample preparation option. DDC-SEM can be successfully applied to any type of soft tissue and cell model where minerals are observed. The sample preparation method of choice has proven invaluable in situations where characterization of pathological minerals is needed in a clinical setting. The sample preparation protocol method of choice has proven invaluable in situations where characterization of pathological minerals is needed in a clinical setting. A fine example is the case of urinary tract stones [12], usually characterized prior to treatment: DDC-SEM can be incorporated in the visualization and characterization of the minerals using the energy dispersive X-ray spectroscopy function of the SEM, which can be applied to provide specific elemental information.

